# Elevated temperatures reduce population-specific transcriptional plasticity in developing lake sturgeon (*Acipenser fulvescens*)

**DOI:** 10.1101/2022.05.28.493847

**Authors:** William S. Bugg, Matt J. Thorstensen, Katie E. Marshall, W. Gary Anderson, Ken M. Jeffries

## Abstract

Rising mean and variance in temperatures elevate threats to endangered freshwater species such as lake sturgeon, *Acipenser fulvescens*. Previous research demonstrated that higher temperatures during development result in physiological consequences for lake sturgeon populations throughout Manitoba, Canada, with alteration of metabolic rate, thermal tolerance, transcriptional responses, growth, and mortality. We acclimated lake sturgeon (30 – 60 days post fertilization, a period of high mortality) from northern and southern populations (56° 02′ 46.5″ N, 96° 54′ 18.6″ W and 50° 17′ 52″ N, 95° 32′ 51″ W respectively, separated by approximately 650 km) within Manitoba to current (summer highs of 20-23^°^C) and future projected (+2-3^°^C) environmental temperatures of 16, 20, and 24^°^C for 30 days, and measured gill transcriptional responses using RNAseq. Transcripts revealed SNPs consistent with genetically distinct populations and transcriptional responses altered by acclimation temperature. There were a higher number of differentially expressed transcripts observed in the southern, compared to the northern, population as temperatures increased, indicating enhanced transcriptional plasticity. Both lake sturgeon populations responded to elevated acclimation temperatures by downregulating the transcription of genes involved in protein synthesis and energy production. Further, there were population-specific thresholds for the downregulation of processes promoting transcriptional plasticity as well as mitochondrial function as the northern population showed decreases at 20^°^C, while this capacity was not diminished until 24^°^C in the southern population. These transcriptional responses highlight the molecular impacts of increasing temperatures for divergent lake sturgeon populations during vulnerable developmental periods and the critical influence of transcriptome plasticity on acclimation capacity.

## 1. Introduction

Freshwater fishes have experienced the highest rates of extinction among all vertebrates throughout the 20^th^ century, influenced by the compounding and dynamic climate change-related stressors of elevated temperatures, hypoxic conditions, and pathogens which continue to imperil freshwater fishes and limit suitable habitat (Marcos-López et al., 2010; Burkhead, 2012; Comte et al., 2012; Schade et al., 2014; Krabbenhoft et al., 2020; Vollset et al., 2020; Jane et al., 2021). Climate change is increasing both the mean and variance of environmental temperature (Thornton et al., 2014; Dillon et al., 2016). Environmental changes are often occurring faster than organisms can adapt to increasing temperatures, especially in species with long generation times (Crozier and Hutchings, 2014; Morgan et al., 2020; Logan and Cox, 2020). Therefore, phenotypic plasticity will be critical for maintaining fitness in response to climate change and promoting survival in these changing environments (Burggren, 2018; Scheiner et al., 2019; Earhart et al., 2022).

Genomic and epigenetic variation can contribute to transgenerational plasticity, while intragenerational plasticity induced to alter an organism’s phenotype at the transcriptomic level may act as an adaptive trait (Muschick et al., 2011; Jonsson and Jonsson, 2019). Together these transgenerational and intragenerational mechanisms provide adaptive plasticity which may enable molecular physiological changes, promoting survival (Lafuente and Beldade, 2019; Oomen and Hutchings, 2017; Wellband and Heath, 2017). Further, polyploid organisms with larger genomes may have increased acclimatory capacity at both the transgenerational and intragenerational timescales through transcriptome plasticity (Ellis et al., 2014; Trifonov et al., 2016; Van de Peer, 2017). In populations of polyploid species with divergent genetic histories, which may be under genetic constraints preventing performance traits from evolving rapidly (Logan and Cox, 2020), genome level changes may be anticipated but alteration of phenotypes through transcriptomic plasticity may be necessary to promote survival as environmental changes intensify.

During early development, change in temperature is a critical abiotic factor necessitating prompt phenotypic alteration. These plastic, individual-level phenotypic changes can alter organismal physiology, buffer against acute environmental changes, lead to long-term phenotypic impacts, and be selected upon across generations (McKenzie et al., 2021). Thus, plasticity and physiological flexibility can be an adaptive response stimulated in stressful environments and is a reversible phenotypic change that may be persistent for days to months, associated with the process of acclimation (Hockachka and Somero, 2002; Crozier and Hutchings 2014; Mackey et al., 2021). Underlying these phenotypic modifications are transcriptional mechanisms that impact the establishment of long-term phenotypes (e.g., developmental plasticity; Jonsson and Jonsson, 2019). Exposing developing fish to increasing environmental temperatures will thus alter traits that may have transient as well as long-lasting physiological consequences. The basis for plasticity is variation in phenotypes produced by the same genotype under different environmental conditions (Lafuente and Beldade, 2019; Earhart et al., 2022). Acclimatory plasticity is especially important in early life to ensure sufficient phenotypic flexibility, enabling fishes to survive in variable environmental conditions until adulthood when their reproductive potential can be reached (Burggren, 2018). In early developmental stages, different phenotypes and developmental trajectories can emerge from genetically distinct populations exposed to similar environmental change based, in part, on differences in transcriptome plasticity (Logan and Cox, 2020). Induced transcriptional responses to stressors often differ between populations and may be indicative of long-term fitness and survival consequences if phenotypes cannot be adequately altered in early life to allow individuals to persist in changing environments (Whitehead et al., 2012; Bugg et al., 2020; Mundy et al., 2020; Bugg et al., 2021).

A primary mechanism underlying physiological responses in the face of environmental change are modifications to the abundance of messenger RNA (mRNA) through transcriptional processes (Smith et al., 2013; Connon et al., 2018; Jeffries et al., 2019; Jeffries et al., 2021). Through mRNA sequencing (i.e., RNAseq) and *de novo* transcriptome assembly, transcriptomic approaches can be used when few genomic resources are available for a species (Alvarez et al., 2015; Connon et al., 2018; Oomen & Hutchings 2017; Thorstensen et al., 2020; Komoroske et al., 2020). Examining genomic sequence differences (SNPs) and differential expression genes through transcriptome-wide approaches can illuminate an organism’s physiologically- and ecologically-relevant responses to environmental change, aid in the identification of tolerance thresholds, and inform conservation management practices (Connon et al., 2018). As environmental extremes in freshwater systems escalate, transcriptomic techniques provide powerful tools to investigate adaptive capacities in species of conservation concern, highlighting the genetic mechanisms that promote physiological acclimation to environmental change. While acclimation may promote thermal resilience, the potential for decreased transcriptional plasticity as well as limitations to other physiological responses in the face of multiple stressors may suggest increased vulnerability, the possibility of severe short-term outcomes, and the potential for impacts on long-term fitness (Smith et al., 2013; Jeffries et al., 2018).

Greater than 50% of the world’s river basins face widespread anthropogenic impacts which limit productivity, species richness, and biodiversity, threatening approximately a quarter of freshwater fish species globally (Palmer et al., 2008; IUCN, 2018; Grill et al., 2019; Su et al., 2021; Warkentin et al., 2022). Habitat loss, commercial exploitation, and changing climates, have pushed many species to the brink of extirpation (Arthington et al., 2016). Sturgeons are some of the most endangered groups of vertebrates on the planet (WWF, 2022), have long generation times (often > 15 years) and have persisted relatively unchanged for hundreds of millions of years. However, because of a rapidly changing environment, sturgeon populations are in widespread decline. Lake sturgeon, *Acipenser fulvescens* are widely distributed throughout North America, and within Canada have experienced population declines >90% in the northern extent of their range since the 1960’s (COSEWIC, 2006). Historically, lake sturgeon population declines in the 1800s were caused by commercial exploitation for caviar. Since the 1950’s however, habitat loss and degradation associated with dams and other barriers have become the most prominent threats to populations and the species’ persistence (Cleator et al., 2010; COSEWIC 2017; Van der Lee and Koops, 2021).

Further, with the intensification of their environments due to changing climates, lake sturgeon now likely face additional stressors, especially so for northern populations. Summer temperatures for Canadian populations can reach upwards of 23^°^C, exceeding temperatures of 20^°^C for approximately 50 days per year, imposing thermal stress and whole-organism physiological impacts for developing lake sturgeon (Bugg et al., 2020). Temperatures in the northern range for lake sturgeon are also increasing faster than the global average (projected 2.1- 3.4^°^C increase by 2050; Manitoba Hydro, 2015), which may further limit suitable habitat, and imperil these endangered sub-Arctic populations as water temperatures exceed their thermal limits (Vincent et al., 2015; Zhang et al., 2019; Bugg et al., 2021).

Increasing environmental temperatures directly influence the development, metabolism, transcriptomic responses, and survival of northern populations of lake sturgeon under laboratory conditions and may contribute to observed declines in wild stocks with high mortality (> 90%) observed in early development (Caroffino et al., 2011; Bugg et al., 2020; Bugg et al., 2021). In fishes like lake sturgeon, the gill lamellae is a multi-purpose tissue comprised of multiple cell types reactive to the external environment with roles in gas exchange, osmoregulation, pathogen defense, acid-base balance, and responses to thermal stress (Lazado and Caipang, 2014; Komoroske et al., 2015; Akbarzadeh et al., 2018; Gilmour and Perry, 2018). Indirectly, elevating temperatures may impact many of these processes in the gill, resulting in increased vulnerability with additional environmental stressors like increase hypoxia or pathogen exposure (LaPatra et al., 1994; Georgiadis et al., 2001; Fujimoto, 2018; Clouthier et al., 2020). Sturgeons may be especially susceptible to the combined effects of increasing temperature, hypoxic environments, and pathogens in early development, when mortality is often high (> 90%; Caroffino et al., 2010) and sturgeon have limited immune responses and ability to escape stressful environments (Webb, 1986; Peake et al., 1997; Schindler, 2001; Deslauriers and Kieffer, 2012; Gradil et al., 2014 a,b; Verhille et al., 2014). However, despite these vulnerabilities in early development, sturgeon have successfully persisted for hundreds of millions of years in ever-changing environments, evolving elaborate polyploid genetic complexity, and demonstrating high levels of phenotypic plasticity in early development, highlighting the potential for inducible transcriptional plasticity (Braasch and Postlethwait, 2012; Trifonov et al., 2016; Bugg et al., 2020; Bugg et al., 2021; Penman, 2021; Yoon et al., 2021; Brandt et al., 2021, Brandt et al., 2022). Therefore, early development represents a crucial window to examine lake sturgeon responses to thermal acclimation, especially within the gill where key homeostatic functions are critical to physiological performance (Burggren, 2018; Barley et al., 2021).

Here we used RNAseq to investigate differential transcriptomic responses to increased environmental temperatures between two latitudinally distinct populations of developing lake sturgeon (McDougall et al., 2017; Bugg et al., 2020). Genomic differences were used to provide context for potential genetic differentiation between the two populations using SNPs in the RNA sequence reads. As sturgeons are ancestral polyploid species, with highly variable genomic architecture and phenotypic plasticity (Braasch and Postlethwait, 2012; Trifonov et al., 2016; Bugg et al., 2021), we expected population-specific responses to temperature treatments in lake sturgeon. We examined the gill mRNA transcript abundance in *n* = 36 individuals from three thermal acclimation treatments in the northern and southern population of lake sturgeon within Manitoba. We targeted early development, 60 days post fertilization, for this analysis to highlight key vulnerabilities to thermal stress for these populations during a critical period in life when high mortality (> 90%; Caroffino et al., 2011) often occurs in the wild. We then investigated phenotypic plasticity through exploration of 9 total contrasts spanning the molecular mechanisms shared between populations as acclimation temperatures increased, as well as population and acclimation-specific responses to increased environmental temperatures. Previous research demonstrated that acclimating lake sturgeon for 30 days to thermally stressful, but ecologically-relevant summer temperatures, of 20^°^C and 24^°^C, resulted in population and acclimation-specific changes in mortality, metabolism, thermal tolerance, and transcriptional responses (Bugg et al. 2020). Therefore, we hypothesized that acclimation to these same elevated temperatures during early life history would elicit transcriptome-wide responses associated with energy mobilization, immune response, cellular repair, and heat shock proteins indicative of chronic thermal stress. Given the higher mortality and lower thermal tolerance observed in the northern population post-temperature acclimation in a previous study (Bugg et al., 2020), we predicted that the northern population of lake sturgeon would show greater transcriptional signatures of thermal stress as temperatures increased, relative to the southern population.

## 2. Materials and Methods

### 2.1 Lake Sturgeon Husbandry and Acclimation

Lake sturgeon used in the current study were from the same populations and acclimation treatments described in Bugg et al. (2020). Briefly, gametes were collected from wild male and female lake sturgeon downstream from the Pointe du Boise Generating Station on the southern Winnipeg River (50^°^17’52’’N, 95^°^32’51’’W) and first rapids on the northern Burntwood River (56^°^ 02’ 46.5’’ N, 96 ^°^54’18.6’’W; separated by approximately 650 km) (Figure 1). Eggs and sperm from the Winnipeg River were fertilized at the University of Manitoba animal holding facility while those from the Burntwood River were fertilized at the Grand Rapids Fish Hatchery, and subsequently transferred to the University of Manitoba. Progeny from the Winnipeg River population were the product of the fertilization of the eggs from two females with the sperm from two males, while individuals from the Burntwood River were the product of the fertilization of eggs from one female with the sperm from six males. Immediately post-fertilization, embryos from each population were submerged in a clay substrate to de-adhease them and then were gently stirred by hand for 1 h. Next, embryos were rinsed with dechlorinated freshwater and kept in tumbling jars to incubate at 12^°^C until hatch. Water temperature was maintained at 12^°^C until 13 days post fertilization after which it was increased by 1^°^C day^−1^ until 16^°^C, matching typical hatchery rearing conditions in Manitoba. Starting at 19 days post fertilization, and continuing throughout the length of the experiment, developing lake sturgeon were fed freshly hatched artemia (Artemia International LLC; Texas, USA) three times daily. Acclimation began at 30 days post fertilization, by increasing treatments to 20^°^C and 24^°^C at a rate of 1^°^C day^-1^ and keeping one treatment from each population at 16^°^C. Lake sturgeon from both populations were acclimated to these temperatures for 30 days with water temperatures recorded by HOBO Water Temperature PROv2 Data Loggers (Onset Computer Corporation; Bourne, MA, USA) and checked by thermometer at least three times daily. All animals in this study were reared and sampled under guidelines established by the Canadian Council for Animal Care and approved by the Animal Care Committee at the University of Manitoba under Protocol #F15-007.

**Figure .**
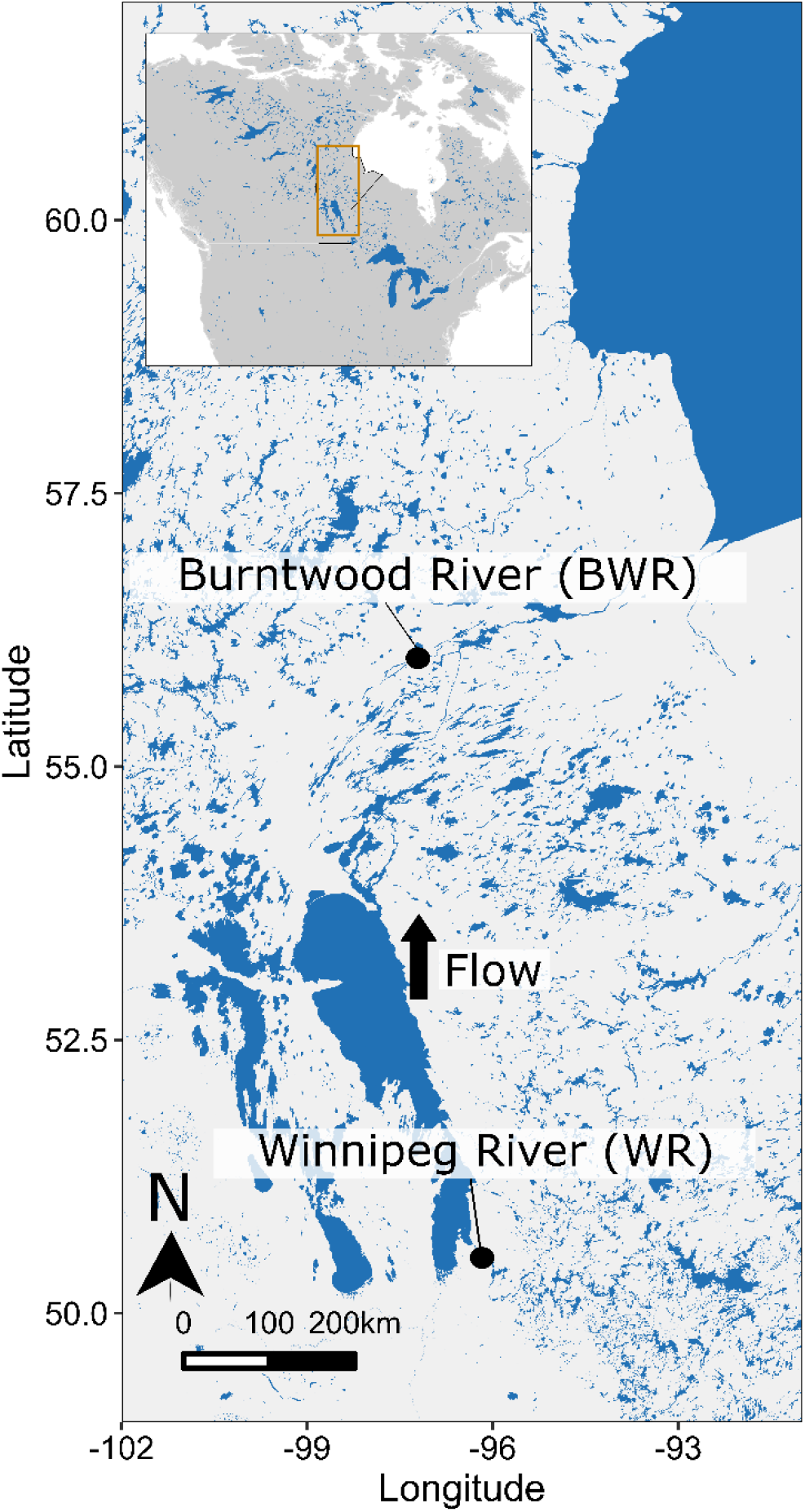
Geographic localities of two lake sturgeon populations (*Acipenser fulvescens*) within Manitoba, Canada, highlighting the northern Burntwood River (BWR) and more southernly (WR) sturgeon populations, separated by approximately 650 km, as well as the direction of flow throughout the watershed. This figure was assembled with the assistance of Evelien de Greef.

**Figure 2.**
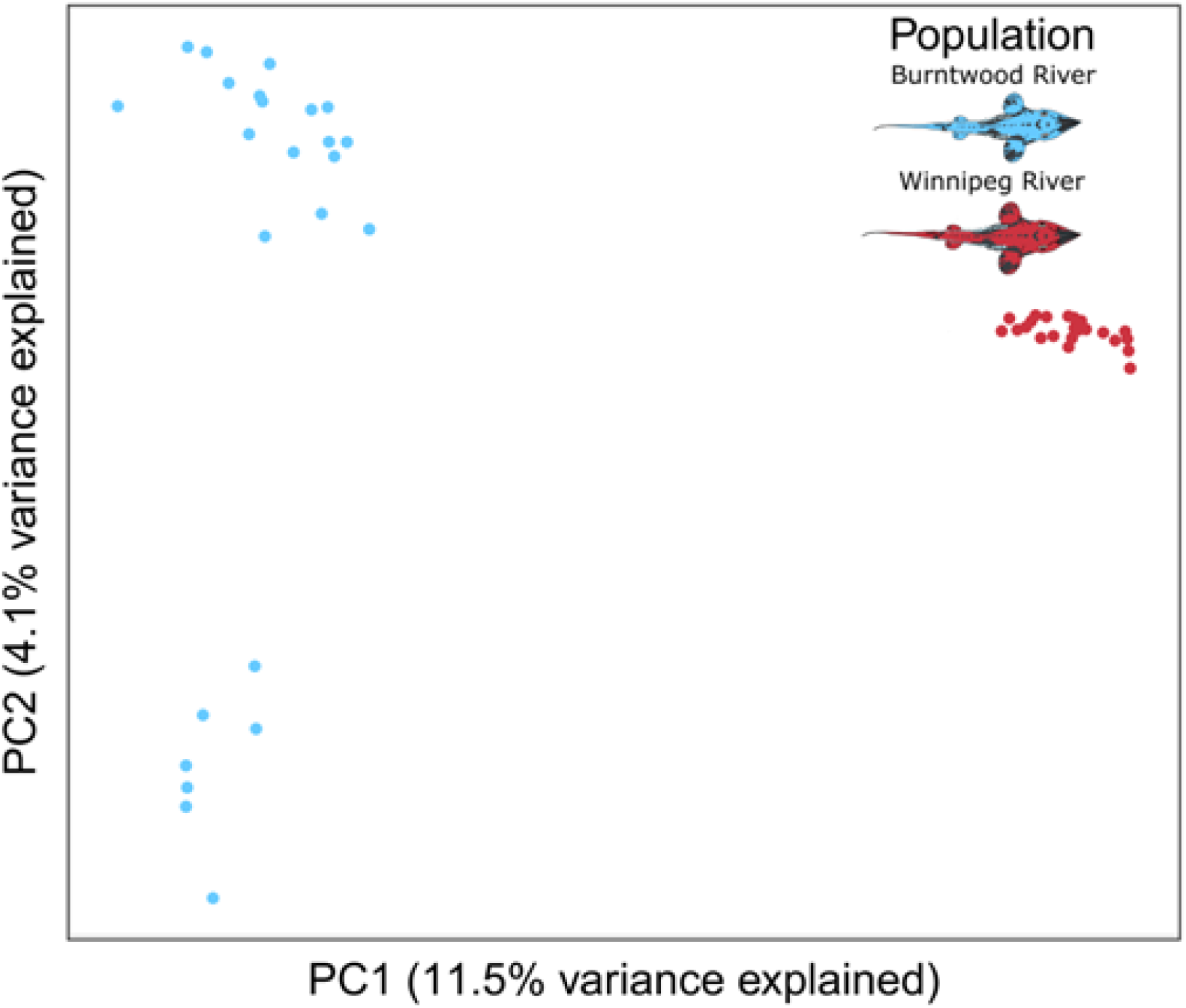
Principal components analysis of 14,517 mRNA SNPs called in lake sturgeon (*Acipenser fulvescens*) from the northern Burntwood River and southern Winnipeg River populations within Manitoba, Canada.

### 2.2 Sampling and RNA extraction

At the end of the 30 day acclimation, six larval lake sturgeon per treatment were netted and euthanized by immersion in an overdose of tricaine methanesulfonate solution (250 mg L^−1^; MS-222, Syndel Laboratory, Vancouver, Canada) with an equal volume of sodium bicarbonate to buffer against pH change. Next, gill tissue was extracted from each larval fish and preserved in RNA*later* (Invitrogen; Carlsbad, CA, USA), held at 4°C for 24 h and stored at −80°C prior to RNA extraction. Total RNA was extracted from the gill tissue by homogenization in 500 µl of lysis buffer (PureLink RNA Mini Kit; Invitrogen; Ambion Life Technologies; Waltham, MA, USA) for 10 min at 50 Hz using a TissueLyser II (Qiagen; Germantown, MD, USA) and then using a PureLink RNA Mini Kit (Invitrogen; Ambion Life Technologies) following manufacturer instructions. Total RNA purity and concentration was evaluated for all samples using a NanoDrop One (Thermo Fisher Scientific) as well as gel electrophoresis to assess RNA integrity. Samples were then stored at −80^°^C prior to sequencing.

### 2.3 *De novo* Transcriptome Assembly

Prior to the present study, a lake sturgeon gill reference transcriptome was created by sequencing mRNA from the pooled RNA of 3 individuals. We used a mRNA isolation, stranded library preparation approach to produce 50 million reads of 100 base pair paired-end reads. The transcriptome was *de novo* assembled using Trinity (Grabherr et al., 2011) through the Canadian Centre for Computational Genomics (C3G; https://computationalgenomics.ca/). Assembled transcripts were then annotated with Trinotate (http://trinotate.sourceforge.net/) using a modified approach. First assembled transcripts were blasted against a fish-specific database with annotation hits returned. Transcripts that were not annotated from the fish specific database were then blasted against the entire SwissProt library.

### 2.4 Sequencing

Total RNA for each sample was normalized to approximately 15 ng μL^-1^ at a total volume of 10 μL and shipped to the McGill Applied Genomics Innovation Core sequencing facility (https://www.mcgillgenomecentre.ca/) for cDNA library preparation and sequencing. Samples were run on a Bioanalyzer to confirm RNA quality and all samples had a RNA Integrity Number (RIN) between 8.7-10. A mRNA isolation step was performed using NEBNext Poly(A) Magnetic Isolation Modules (New England Biolabs) followed by using NEBNext Ultra II Directional RNA Library Prep Kit for Illumina (New England Biolabs) to produce stranded cDNA libraries. NEBNext dual adaptors (New England Biolabs) were then individually applied to barcode each library prior to sequencing of 100 base pair paired-end reads. All 36 sturgeon gill samples were sequenced on a single NovaSeq 6000 (Illumina) lane producing 1.53 billion total reads with an average of 42,528,434 (± 17,812,372 SD) reads per sample.

### 2.5 Quality control and Trimming

Raw read files were uploaded to Galaxy servers (Giardine et al., 2005) and paired reads were joined and checked for quality using FastQC version 0.11.8 (Andrews, 2010). To filter low-quality reads, Trimmomatic version 0.38.0 was applied with the Illuminaclip option set to standard for TruSeq3 adapter sequences, a maximum mismatch count of 2, and a palindrome read alignment threshold of 30. Additionally, a sliding window trimming step was used with a size of 4 base pairs as well as a minimum read length filter set at 20 bases. Ultimately this trimming removed approximately 2.8% (± 0.26% SD) of reads from each file. Full details for quality control and trimming procedures are available in Supplementary File 1, Sheets A-C.

### 2.6 Read Mapping and Differential Expression

After filtering, HISAT2 version 2.1 and StringTie version 2.1.1 (Kim et al., 2015; Pertea et al., 2015) were used to align paired read libraries to the *de novo* assembled gill transcriptome and produce a set of assembled transcripts for each sample, respectively. HISAT2 was run with default parameters and produced an overall alignment rate of approximately 83% (± 0.7% SD) per sample. HISAT2 aligned bam files were then input into StringTie to assemble transcripts, using default settings, producing an average of 94,992 (± 11,075 SD) transcripts per sample. StringTie files for each sample were then merged producing a master list of 139,456 transcripts.

To assess differential transcript expression each of these transcripts were first counted in HISAT2 produced aligned read files using featureCounts version 1.6.2 with default settings (Liao et al., 2014; Available in Supplementary File 2). Total counts were then assessed for differential expression using the R v4.0.0 package EdgeR v3.32.1 (Robinson et al., 2010; R Core Team, 2022). Prior to analysis the “filterByExpr” function with default arguments was used to retain transcripts expressed among any individual, resulting in a final count of 95,593 transcripts. Full details for read mapping of each individual sample are available in Supplementary File 1, Sheet D.

### 2.7 Principal Component Analysis, Differentially Expressed Transcripts, and Aggregate Heatmapping

A principal component analysis (PCA) was conducted on featureCounts from the above 95,693 transcripts after normalizing the data to the library size using the plotMDS function in the limma package (Ritchie et al., 2015). Next, numbers of differentially expressed genes between the Burntwood River and Winnipeg River populations of lake sturgeon acclimated to 16^°^C, 20^°^C, and 24^°^C for 30 days in early development were tabulated for upregulated and downregulated genes independently, representing genes with a log_2_ fold change > 1 and a *q*-value (FDR adjusted p value) < 0.05 for each of the 9 individual contrast (Details for all differentially expressed transcripts as outlined in Figure 3B can be found in Supplementary File 3, Sheets A-C). The 9 total contrasts compared the populations at each acclimation temperature as well as across acclimation temperatures for each population. Log_2_ counts per million (CPM) data for all 95,693 transcripts were then generated and the top 500 genes from each contrast with the highest absolute value log_2_ CPM fold change were included in an aggregate list of the most highly differentially expressed genes across all contrasts, resulting in a total of 4,500 transcripts, 2,285 uniquely included (full CPM data are available for all 95,693 transcripts in Supplementary File 4). A heatmap with hierarchical clustering was then produced using the log_2_ CPM expression of these aggregate most highly differentially expressed 2,800 unique transcripts using the pheatmap package (Kolde, 2015).

**Figure 3.**
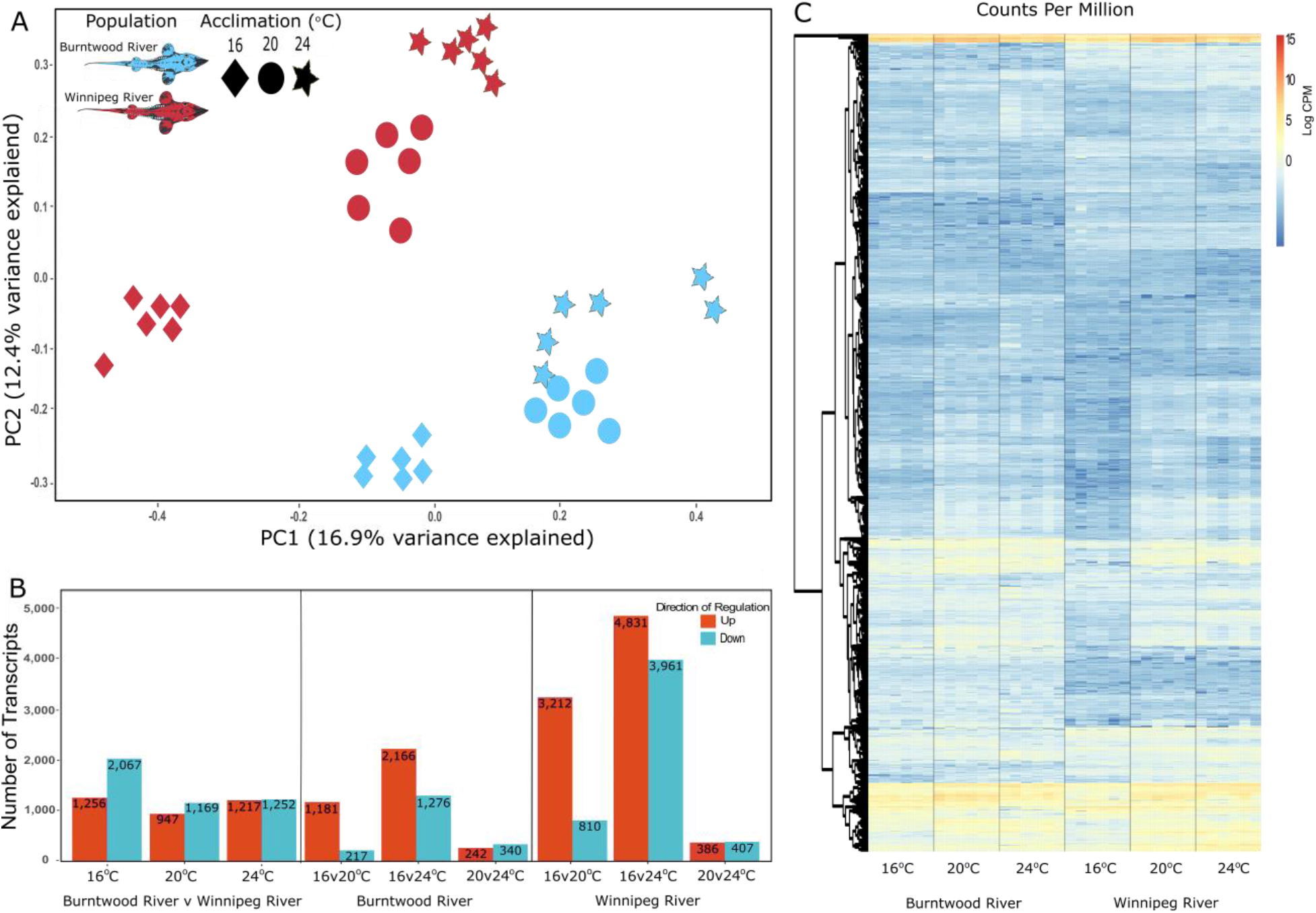
A) Principal component analysis and B) Numbers of differentially expressed transcripts across all transcripts, as well as C) Aggregate heat map of the top 500 differentially expressed transcripts between each contrast of northern Burntwood River and southern Winnipeg River populations of lake sturgeon within Manitoba, Canada, acclimated to 16^°^C, 20^°^C, and 24^°^C for 30 days in early development, measured using RNAseq. Transcript numbers represent upregulation and downregulation with a log_2_ fold change > 1, respectively, and with a *q*-value (FDR adjusted p value) < 0.05. Contrasts are oriented Burntwood River-Winnipeg River at the interpopulation level and lower temperature-higher temperature across acclimation treatments (e.g. up means increased with temperature, and in the Burntwood River population, relative to Winnipeg River). Each population and acclimation treatment is represented by 6 individual gill samples (n = 6). For the aggregate heat map, only unique transcripts were included for a total of 2,800 transcripts represented. Data are expressed as log_2_ counts per million (CPM).

### 2.8 Functional Term Enrichment

Transcripts annotated to the *de novo* lake sturgeon transcriptome were split into groups of upregulated and downregulated genes by treatment based on their log_2_ CPM fold change (< −1 or > 1) and were counted for distinct gene names (hereafter referred to as split-contrast). The positively and negatively regulated annotated genes were then analyzed for functional enrichment using gene ontology (GO) terms through enrichR (Chen et al., 2013; Kuleshov et al., 2016; Xie et al., 2021) with positive hits returned from the “GO_Biological_Processes_2018”, “GO_Molecular_Function_2018”, and “GO_Cellular_Component_2018 databases and filtered for an adjusted *p*-value < 0.05. The resulting 213 enrichment terms across the upregulated and downregulated spit-contrast were then tabulated into a master file detailing their presence or absence in each split-contrast (0 or 1; Full presence and absence data across each split-contrast is available in Supplementary File 5). This master file was then used to create UpSet plots of the shared and population-specific terms both upregulated and downregulated across all split-contrasts, upregulated terms as temperatures increased from 16^°^C to 20^°^C, and terms which decreased as temperatures increased from 20^°^C to 24^°^C using the ‘UpsetR’ package v1.4.0 (Conway et al., 2017). As the split-contrasts with the most GO terms were largely downregulated, split-contrasts with downregulated functional enrichment terms as temperatures increased were then focused on for interpretation of results.

Next, non-redundant enrichment terms in the GO Biological Process category were identified with REVIGO for summary and visualization (Supek et al., 2011). Adjusted *p* values were supplied for all REVIGO analyses to guide the clustering of redundant GO terms by the REVIGO algorithm (Supek et al., 2011). Where GO terms were shared across populations or acclimation treatments, the geometric mean of the adjusted *p* values from all shared groups was input into REVIGO. All REVIGO output is available in Supplementary File 6, Sheets A-I.

### 2.9 Aggregate Gene Network Analysis and Enrichment

An aggregate of the most highly differentially expressed transcripts was then investigated, producing a total of 503 uniquely annotated most differentially expressed transcripts, from the aggregate list of 4,500 most differentially expressed transcripts across contrasts used for the heatmap (the full aggregate list of 503 annotated most differentially expressed transcripts is available in Supplementary File 7). These 503 unique gene names were uploaded to OmicsNet (Zhou and Xia, 2018) and input under the genes category using the “official gene symbol” classification and zebrafish *Danio rerio* as the study organism, as this was the most closely related model species option (used for OmicsNet aggregate network analysis only). Next, protein-protein interactions were selected in the network building panel, using STRING the database (a comprehensive collection of physical and functional protein-protein interactions covering 5090 organism across a broad and diverse selection of benchmarked data sources; Szklarczyk et al., 2019), to assembly a network of the unique genes and their potential interactions. The network was then changed to a 2-D model, using a force atlas layout, with seed nodes highlighted with a blue ring around the node, and edge bundling applied. Functional analysis of the produced network was then categorized using the Function Explorer, querying all nodes against the GO: Biological Process database. Processes with a *p* value < 0.05 are reported in Supplementary File 8. Additionally, degree of connectivity and node betweenness scores were tabulated for each transcript in the network and are included in Supplementary File 8 as well.

### 2.10 SNP Calling and Analyses

Population differentiation was assessed with a combination of RNA-seq and reduced representation-oriented programs to mitigate issues with population genetic analyses of polyploid mRNA data (Thorstensen et al., 2021; Thorstensen et al., 2022a). The program fastp v0.20.1 was used for adapter trimming and read filtering, with a minimum phred quality of 15, a maximum of 40% of unqualified bases allowed in a read before read filtering, a minimum length of 100 base pairs, and polyG read tails were trimmed (Chen et al., 2018). STAR v2.7.9a was used as a splice-aware aligner in two-pass mode to map reads to the lake sturgeon gill transcriptome used for differential expression analyses in this study (Dobin et al., 2013; Thorstensen et al., 2022a).

Picard v2.26.3 was used to clean the resulting bam files, add read groups, and mark duplicate reads. GATK v4.2.4.0 was used to split reads that contained Ns in cigar reads (Van der Auwera et al., 2013). Next, STACKS v2.60 was used to create loci for SNP calling (gstacks) and call SNPs (populations) with a minimum minor allele count of 2, at least 60% of the data present within each of the Winnipeg and Burntwood River groups, and 90% present data overall (Catchen et al., 2013). Because paralogous loci may lead to biased population genetics measures, HDplot was used to filter out loci with heterozygosity ≤ 0.7 and read ratio deviations ≤ 7 (McKinney et al., 2017). SNPRelate v1.30.1 was used to filter out SNPs with R^2^> 0.20 along 500 kilobase segments, treating each transcript as a chromosome (Zheng et al., 2012).

Hierfstat v0.5-11 was used to assess Weir & Cockerham’s *F*_ST_ between the samples from the Burntwood and Winnipeg Rivers, with 95% confidence intervals generated by 1000 bootstrap iterations (Goudet, 2005). Admixture v1.3.0 was used to estimate ancestry coefficients for each group between K values of 2, and pophelper v2.3.1 was used to visualize the results from Admixture (Alexander et al., 2009; Francis et al., 2017). While further population genetic and association-based analyses were possible (e.g., tests for signatures of selection), extensive family structure remained in the data because genotyped samples were *not* random representations of their populations of origin. Therefore, the present population genetic analyses were used to assess genetic dissimilarities between groups in the context of other research and to demonstrate how transcriptional differences between groups may be consistent with underlying genetic differences.

## 3. Results

### 3.1 Principal Component analysis, Differentially Expressed Transcripts and Aggregate Heatmapping

Examination of the PCA (Figure 3A), number of upregulated and downregulated genes (Figure 3B), as well as the heatmap displaying the most highly differentially expressed genes across contrasts (Figure 3C), demonstrated that the Burntwood River and Winnipeg River populations of lake sturgeon responded differently to elevated temperatures both in the magnitude and trajectory of transcriptional change. The PCA showed divergent response patterns with Winnipeg River sturgeon in treatments with populations grouping out separately along a linear trajectory as temperatures increased. Acclimation treatments primarily separated out linearly across principal component 1 (16.9% contribution), while populations were dispersed vertically across principal component 2 (12.4% contribution). As temperatures increased from 16^°^C to 20^°^C both higher temperature treatments separated out cleanly from their 16^°^C counterparts. However, as temperatures increased from 20^°^C to 24^°^C, these two temperature treatments remained with some overlap for the northern Burntwood River population, while the 24^°^C treatment for the southern Winnipeg River population continued on its responsive trajectory, separating out from the 20^°^C treatment.

Patterns of upregulation and downregulation between the two populations at a given acclimation temperature, and within populations across acclimation temperatures, demonstrated population-specific and acclimation-specific responses to elevated temperatures in developing lake sturgeon. When comparing Burntwood River and Winnipeg River sturgeon at each acclimation temperature there are similar numbers of up and downregulated transcripts. However, within population comparisons reveal a much higher level of transcriptional plasticity in the southern Winnipeg River population. The largest changes in transcript regulation in the Burntwood River and Winnipeg River sturgeon occurred as acclimation temperatures increased from 16^°^C to 24^°^C, where the populations differentially expressed a total of 3,442 and 8,792 transcripts, respectively. These changes in transcript expression between 16^°^C and 24^°^C represent the differential expression of approximately 3.6% and 9.2% of all transcripts for the Burntwood River and Winnipeg River populations of sturgeon included in the analysis, respectively. Between 16^°^C and 20^°^C these numbers of differentially expressed transcripts were less than half when compared to the 16 to 24^°^C transition, with just 1,398 and 4,022 transcripts differentially expressed in the Burntwood River and Winnipeg River, respectively. While producing similar qualitative patterns of expression to the Burntwood River, the number of differentially expressed transcripts was higher in the southern Winnipeg River across acclimation treatments, with lower values in the northern latitude Burntwood River population.

The heatmap outlined specific patterns of transcript expression of the most highly differentially expressed transcripts, with divergent log_2_ CPM expression values as temperatures increased above 16^°^C for both populations but with differences in these patterns across populations (Figure 3C). Differential expression patterns of these transcripts are largely qualitatively similar across the Burntwood River 20^°^C and 24^°^C treatments, with those from the Winnipeg River 20^°^C and 24^°^C treatments. However, there is evidence of a much larger change in transcriptional regulation observed between the Winnipeg River 16^°^C and higher temperature treatments (20^°^C and 24^°^C), especially when compared to patterns of expression observed in the Burntwood River sturgeon as temperatures increase. Overall, the patterns in the most highly differentially expressed genes align with the PCA in the increased separation between Winnipeg River 16 and 20^°^C acclimation treatments, and the lack of separation between Burntwood River 20 and 24^°^C acclimation treatments compared to Winnipeg River sturgeon in the same temperatures. Overall, these patterns suggest that the most highly differentially expressed transcripts reflect similar transcriptional trajectories as shown in the PCA of all transcripts.

### 3.2 Functional Term Enrichment

Term enrichment demonstrated 213 unique GO pathway terms distributed across the 17 split-contrasts (Supplemental Figure 2A). Overall, the split-contrasts with the largest number of assigned terms were those underlying processes being upregulated as temperatures increased from 16 to 20^°^C (WR16v20_neg and BWR16v20_neg; 75 and 85 terms respectively) and those processes that decreased in expression as temperatures continued to increase from 20^°^C to 24^°^C (WR20v24_pos and BWR20v24_pos; 62 and 97 terms, respectively). The split-contrasts between the Burntwood River and Winnipeg River populations as temperatures increase from 16^°^C to 20^°^C and again from 20^°^C to 24^°^C specifically revealed conserved and population-specific biological processes upregulated as temperatures increase from 16^°^C to 20^°^C and those that are decreased with further temperature increase to 24^°^C. As temperatures increased from 16^°^C to 20^°^C there were 50 upregulated functional term responses shared between populations, as well as 39 responses specific to the northern Burntwood River population and 25 distinct to the southern Winnipeg River population (Supplemental Figure 2B). As acclimation temperatures increased further from 20^°^C to 24^°^C there were 26 functional term responses shared between populations, with 71 terms specific to the Burntwood River population and 38 distinct to the Winnipeg River population (Supplemental Figure 2B).

### 3.3 GO Term and REVIGO Analysis

Overwhelmingly, the majority of all 213 terms were categorized as GO Biological Processes, encompassing 77% of terms identified and highlighting transcriptional mechanisms altered across lake sturgeon populations as temperatures increase, therefore these terms were focused on for further analysis.

#### Conserved responses to increasing temperature

Across both temperature increases (16 to 20^°^C and 16 to 24^°^C), there were 8 GO Biological Process terms identified as downregulated and conserved across both populations. These mechanisms were involved in RNA maturation, mRNA splicing, mitochondrial protein insertion, and ribosome biogenesis (Figure 4; Supplementary File 6, Sheet A).

**Figure 4.**
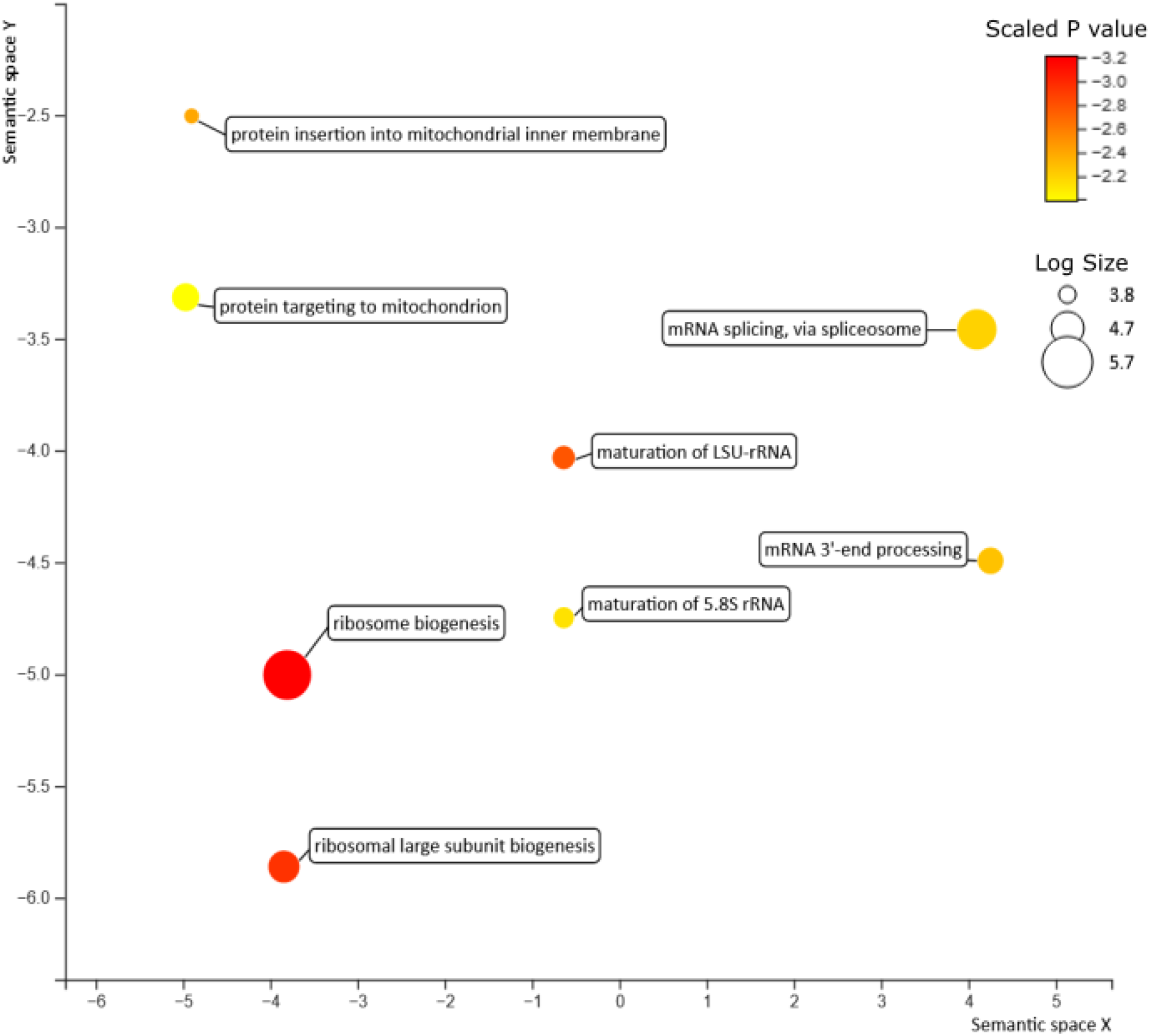
REVIGO (Reduce + Visualize Gene Ontology) plots of GO (gene ontology) biological processes terms that decreased during a 30-day thermal acclimation from both 16 to 20°C and 16 to 24°C for developing northern and southern lake sturgeon (*Acipenser fulvescens*) populations within Manitoba, Canada, measured using RNAseq. These terms were conserved across both the southern Winnipeg River (WR) and northern Burntwood River (BWR) lake sturgeon populations as temperatures increased. Bubble color indicates the significance of the adjusted *p* value at after applying a log_10_ scale (darker color is a more significant term), while bubble size indicates the frequency of a GO term GO annotation database (larger bubble is a more common GO term in the data set).

As the acclimation temperature increased from 16^°^C to 20^°^C there were 29 conserved GO Biological Process terms identified as downregulated and shared between the northern Burntwood River and southern Winnipeg River populations of lake sturgeon. These shared processes were involved in chromosomal structure (telomere maintenance, histone methylation), mRNA processing and maturation (maturation of rRNA, mRNA splicing, mRNA processing, miRNA-mediated gene silencing), ribosomal biogenesis and protein modification (ribosome biogenesis, protein methylation), as well as mitochondrial function (protein targeting to mitochondrion, protein insertion into mitochondrial inner membrane; Supplementary File 6, Sheet B).

As acclimation temperature further increased from 16^°^C to 24^°^C there were 22 conserved GO Biological Process terms that decreased in expression in both the Burntwood River and Winnipeg River lake sturgeon populations. These shared processes were involved nuclear regulation and DNA repair (double-strand break repair, mitotic nuclear membrane reassembly, mitotic nuclear membrane organization, DNA duplex unwinding), further mitochondrial dysregulation (mitochondrial translation, elongation, organization, protein localization, in addition to terms above), mRNA and RNA splicing (via spliceosome, and transesterification reactions respectively), ribosomal biogenesis, and protein insertion into the ER membrane (Supplementary File 6, Sheet C), demonstrating further disruption at the DNA, mRNA, mitochondrial, and protein synthesis levels as temperatures increased.

#### Population-specific responses to increasing temperature

As temperatures increased from 16^°^C to 20^°^C, there were also population-specific increases in transcripts involved in various biological processes, fairly evenly split across both populations. Specific to the Winnipeg River population, there were 38 GO Biological Process terms identified as downregulated (Supplementary Figure 3A) largely characterized by changes in mRNA modification and transcription (gene expression, mRNA processing, non-sense mediated decay, tRNA export, mRNA stability regulation, RNA polymerase I transcription, mRNA cleavage, RNA mediated gene silencing), protein production and modification (ribosomal disassembly, translation, regulation of translation, translation termination, peptide biosynthetic process, amino acid modification), signaling (negative regulation of Wnt signaling pathway, tyrosine phosphate signaling pathway, G1 and p53 DNA damage signaling), and mitochondrial function (mitochondrial gene expression, protein localization, as well as translational elongation and termination; Supplementary File 6, Sheet D). In the Burntwood River population, there were an additional 20 population-specific GO Biological Process terms downregulated (Supplementary Figure 3B). These Burntwood River specific upregulated terms involved further impacts on mRNA processes and alternative splicing mechanisms (histone arginine methylation, peptidyl-lysine monomethylation, regulation of alternative splicing via spliceosome, mRNA 3’end processing, RNA nuclear export, miRNA metabolism, and different signaling mechanism from the Winnipeg River population (positive regulation of Wnt signaling, canonical Wnt signaling pathway, smoothened signaling pathway, regulation of p38MAPK cascade; Supplementary File 6, Sheet E). Additionally, the Burntwood River population exhibited population-specific mitochondrial processes terms including lipid metabolism and mitochondrial respiratory chain complex assembly, key for energy production.

As temperatures increased further from 16^°^C to 24^°^C there were more population-specific GO Biological Process terms identified as downregulated, however these changes were almost solely in the Winnipeg River population, with 56 population-specific process to 3 in the Burntwood River population (Supplementary Figure 3C). These terms involved increased signs of cellular disfunction with downregulation of processes involving translational disruption, telomere maintenance (via telomere lengthening and telomerases) DNA damage (cellular response to DNA damage, nucleotide excision repair, DNA repair), mitochondrial and oxidative stress (respiratory chain complex III assembly), ubiquitin and protein cycling (ubiquitin protein transferase and ligase activity), and immune function (regulation of type I interferon signaling; Supplementary File 6, Sheet F). In contrast, there were only 3 population specific processes for this temperature contrast in the Burntwood River involving the downregulation of endosomal transport via the multivesicular body sorting pathway, neuromuscular processes, and ureter development.

### 3.4 Aggregate Transcript Network Analysis and Functional Enrichment

The aggregate most highly differentially expressed transcript network demonstrated connections across many different functional groups of transcripts that were altered in developing lake sturgeon during acclimation to elevated temperatures. Transcripts identified as hubs (orange circles, highlighted in blue; Figure 5) form many responsive network connections and are active in DNA regulation and repair processes (*cdk2*, *ddit4, trip12, exo1, mcm9, chd1l, smarcad1a*), mRNA splicing (*zmat5, cwc27, prpf31*), protein turnover and synthesis (*rps27a, cops5, hap1, rnf8, usp13, fbxo45, smurf2, rplp0, bop1, eif3d*), chaperone activity and protein folding (*hsc70, hspa8, hyou1, dnajc2, cct4, unc45b, naca*), as well as immune and inflammatory activation (*traf6, b2m, banf1, rsad2, egr1, jak1, prkcbb*). Additional hub genes were involved in more general processes like calcium regulation (*fyna, cam2kd1*), cell cycling (*plk4, spc24, cdc14ab, pds5a*), autophagy (*atg4b, trim13*), development (*kif7*), osmotic balance (*cftr*) and mitochondrial function (*gfm1*). Many of these processes were also present in the network GO Biological Process term enrichment with enriched pathways including protein ubiquitination, catabolism, folding and stability, RNA splicing (including spliceosomal assembly), immune function (inflammation, virus, immune system process, adaptive immune response, cytokine-mediated signaling pathway, lymphocyte differentiation), responses to DNA damage and repair, heart development, and responses to stress (including heat; Supplementary File 8). The hub transcript with the highest degree of connectivity and network betweenness score (Supplementary File 8) was *rps27a* with dual roles in ribosome and ubiquitin structure.

**Figure 5.**
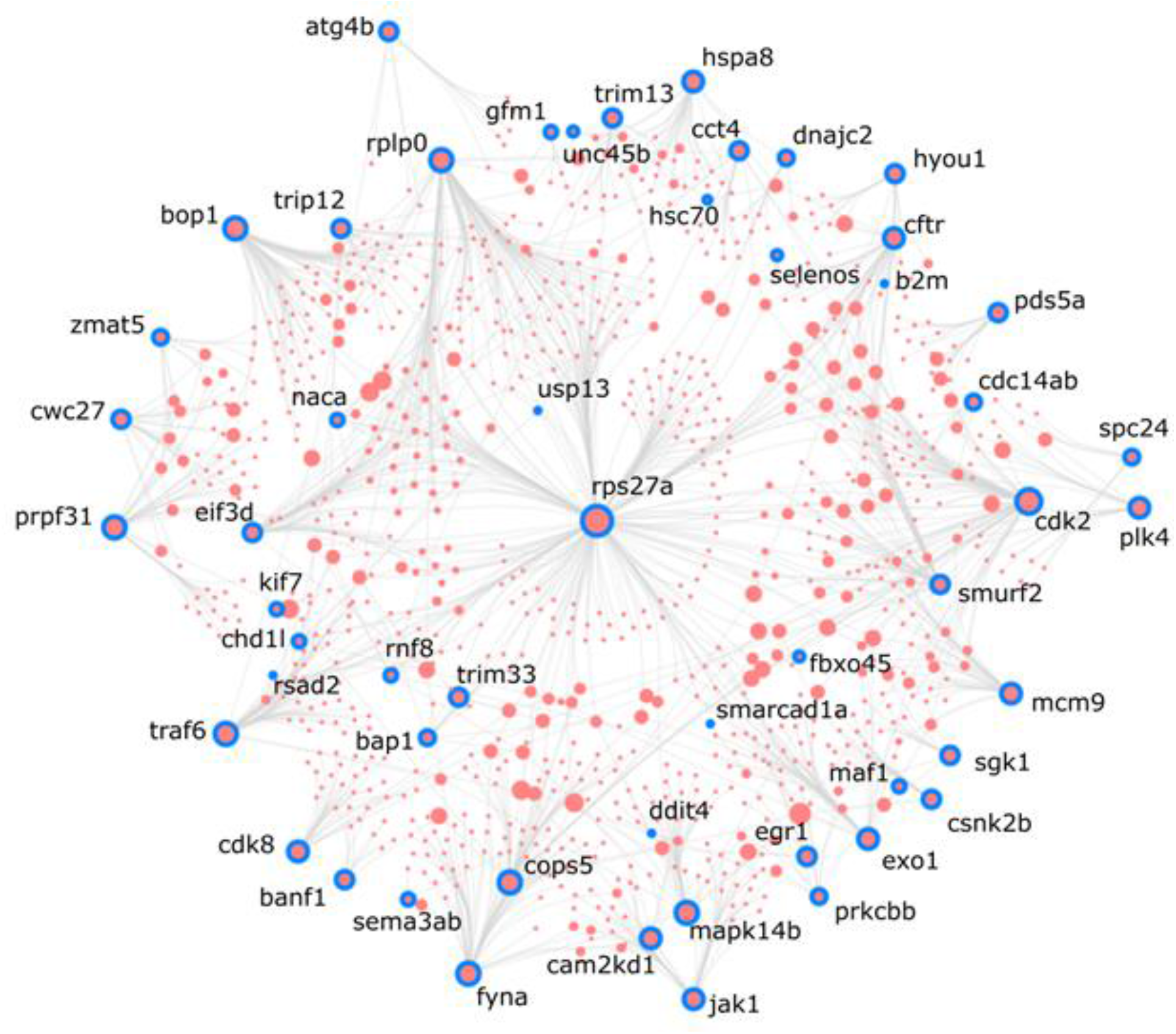
Aggregate most differentially expressed transcript network demonstrating predicted protein-protein interactions across the most upregulated and downregulated transcripts of northern and southern lake sturgeon, *Acipenser fulvescens*, populations, within Manitoba, Canada, acclimated to 16^°^C, 20^°^C, and 24^°^C for 30 days in early development, measured using RNAseq. Orange transcript circles outlined in blue with bolded names indicate hub transcripts within the network. Hub size and naming is scaled according to their degree of interaction (number of connections) with surrounding hubs (Zhou and Xia, 2019).

### 3.5 SNP Calling and Analyses

An initial 32,244 SNPs were called with STACKS from the mapped reads, of which 24,396 were accepted as non-paralogous, and 14,517 were accepted after pruning for linkage disequilibrium. This dataset of 14,517 SNPs was used for subsequent analyses.

A principal component analysis for SNPs between lake sturgeon samples in the present data demonstrate that the northern Burntwood River and southern Winnipeg River populations are genetically distinct (Figure 2) with an *F*_ST_ of 0.086 (0.082 – 0.090 95% CI). The admixture plot (Supplemental Figure 1) demonstrates distinct populations with ancestry from the southern Winnipeg River population in the downstream northern Burntwood River population attributable to introgression or uncertainty in ancestry coefficient estimates.

## 4. Discussion

In the current study, we demonstrated conserved and population-specific transcriptional responses as environmental temperatures increased in two geographically and genetically distinct endangered populations of lake sturgeon. These responses spanned wide ranging cellular protective processes induced during acclimation to thermally stressful conditions. Specifically, these processes related to the cellular machinery involved in transcriptional and translational regulation, mitochondrial function, oxidative damage, and immunocompetence. Overarching transcript expression patterns indicated population-specific capacity for gene transcription as environmental temperatures increased, with divergent transcript response trajectories, different temperature thresholds to induce transcriptional change, and a higher magnitude of transcriptional plasticity apparent in the southern Winnipeg River population. Conserved and population-specific biological process responses illustrated that this transcriptional plasticity promoted whole-organism acclimation to elevated temperatures in developing lake sturgeon with potential for adaptive responses at the population scale. Ultimately, these processes contribute to the ability of lake sturgeon to acclimate to environments with increasing temperature and highlight the physiological consequences once acclimatory capacity has been exceeded.

### Plasticity as an Adaptive Trait

Thermally responsive molecular pathways first rely on the ability of lake sturgeon to transcriptionally respond to environmental change. Without the necessary translational machinery in place, mRNA abundance changes cannot effectively modify the proteome or the physiology of the organism (Vornanen et al., 2005; Smith et al., 2013; Akbarzadeh et al., 2018). Thus, as thermal thresholds are reached and lake sturgeon face a reduction in the ability to transcribe mRNA’s, produce ribosomes, and maintain mitochondria, those consequences likely negatively impact the organism’s ability to respond to changes in their environment with impaired and diminished mRNA, protein, and energy production.

Increasing temperatures induced alteration in the abundance of transcripts associated with post-transcriptional modifications in both populations of lake sturgeon, influencing cellular plasticity. Conserved responses of alternative splicing and gene silencing via miRNA suggest that these mechanisms are part of the inducible stress response of sturgeons and likely play a role in regulating phenotypic plasticity, as observed in other fishes (Somero, 2018; Healy & Schulte, 2019; Tan et al., 2019; Shiina and Shimizu, 2020; Verta and Jacobs, 2021; Thorstensen et al., 2022b; Steward et al., 2022). However, these processes were ultimately downregulated as temperatures increased to 24^°^C, indicating thermal thresholds on post-translational as well as transcriptional responsiveness. The conservation of these mechanisms across both populations of lake sturgeon, as temperatures increase and under stressful conditions in various other species of fish, highlight the importance of post-transcriptional modifications in response to environmental stressors across the evolution of fish lineages.

As temperatures surpass thermal thresholds, and the effectiveness of some transcriptional and translational mechanisms are diminished, overall physiological plasticity is likely reduced. Fishes’ susceptibility to the effects of additional stressors are thereby likely increased by compromising their ability to respond at multiple levels of biological organization (Smith et al., 2013; Brennan et al., 2022). As temperatures increased, transcription modifying processes were more highly preserved in Winnipeg River sturgeon that historically experience higher temperature thermal environments in the wild. As the fish used from each wild population are genetically distinct (*F*_ST_ of 0.086), experience different thermal environments, and have been geographically separated throughout recent history (McDougall et al., 2017; Bugg et al., 2020), the transgenerational preservation of these transcriptional mechanisms may itself be an adaptive mechanism, enhancing the southern Winnipeg River population’s plasticity and ability to respond to environmental change (Fox et al., 2019). Enhanced transcriptional plasticity likely results in the observed increases in growth, metabolism, thermal tolerance, and survival of the southern Winnipeg River population under thermally stressful conditions, when compared to their Burntwood River counterparts (Bugg et al., 2020).

### Transcriptomic Responses to Increasing Temperatures

As temperatures increased first from 16^°^C to 20^°^C and then to 24^°^C, analysis of downregulated biological processes revealed 8 shared transcriptional responses conserved between the lake sturgeon populations. Together, these processes illustrated the effects of increased temperatures with impacts on key cellular mechanisms involved in transcription, translation, and energy production, processes fundamental to cellular function. Without the necessary cellular machinery in place, the capacity for protein level changes to plastically alter the phenotype of developing lake sturgeon and promote environmental acclimation at the whole organism level is diminished (Buccitelli and Selbach, 2020). Further, acclimation to 20^°^C was associated with decreases in processes involved in telomere organization in both populations, which have the potential to disrupt normal cell cycling and proliferation (Barrington et al., 2017). In both populations, temperature increase to 24^°^C highlighted decreases in DNA repair mechanisms as thermal stress mounts, which can result in a wide array of cellular consequences such as genome instability, modulation of disease-states, impacts to cellular health and ultimately senescence (Chatterjee and Walker, 2017; Barnes et al., 2019). These impacts of temperature increase likely elevate the lake sturgeon’s susceptibility to the effects of compounding environmental stressors, which may be expected as environmental conditions intensify (Smith et al., 2013; Todgham and Stillman, 2013; Brennan et al., 2022).

Population-specific responses to increased temperature highlight transcriptional mechanisms specific to the northern or southern lake sturgeon. As temperatures increased to 20^°^C, the northern Burntwood River population decreased processes promoting transcriptional plasticity including alternative splicing and methylation (similarly as observed in Atlantic Sturgeon, *A. oxyrinchus;* Mastrangelo et al., 2012; Metzger and Schulte, 2016; Steward et al., 2022; Thorstensen et al., 2022b; Penny et al., 2023), as well as mitochondrial processes involved in energy production such as lipid metabolism and respiratory chain complex assembly (Greene and Selivonchick 1987, Chung and Schulte, 2015). In contrast, the southern Winnipeg River population demonstrated decreases in cellular processes dominated by mRNA modification and transcriptional mechanisms. For the Winnipeg River population, downregulation of genes involved in alternative splicing and methylation, and energy producing mechanisms did not occur until 24^°^C was reached, highlighting potential mechanisms that underly observed population-specific thresholds in transcriptional plasticity as well as mitochondrial function and energy production (Bugg et al., 2020). Overall, as temperatures reached 24^°^C population-specific processes primarily occurred only in the southern Winnipeg River population, further demonstrating the preservation of transcriptional plasticity in this population. Specifically, we observed downregulation of mechanisms involved in response to DNA damage, oxidative stress, protein turnover, and immune function. As the southern Winnipeg River population is exposed to higher yearly temperatures and demonstrates population-specific transcriptional processes when exposed to increased temperatures, as compared to their more northerly Burntwood River counterparts (Bugg et al., 2020; Bugg et al., 2021), these unique responses are likely indicative of adaptive mechanisms downregulated to preserve energy and promote survival as temperatures increase (Fox et al., 2019).

### Network Analysis

Predictive transcript network analysis, of the aggregate most highly differentially expressed transcripts, revealed functional processes possibly significant for thermal acclimation. Key hub transcripts and enriched pathways in these regulatory networks involved impacts on DNA integrity, mRNA splicing, protein turnover and synthesis, chaperone activity and protein folding as well as immune responsive mechanisms, with within-network GO analysis indicating the recruitment of similar biological processes. Together, these highlighted transcripts and pathways indicate the importance of these systems in the transcriptomic response to thermal acclimation across different populations of developing lake sturgeon.

The effects of thermal stress had direct impacts on genes involved in the integrity of DNA, altering damage signaling, repair processes, and cellular cycling (pathways: DNA repair, response to DNA damage stimulus, DNA damage checkpoint; hub transcripts: *cdk2*, *ddit4, trip12, exo1, mcm9, chd1l, smarcad1a*). These transcripts are largely involved in damage detection during the G1/S phase transition, and associated DNA replication (Bacevic, et al., 2017) suggesting key cellular processes disrupted by thermal stress in lake sturgeon. This G1 checkpoint specifically detects DNA damage prior to entering the S phase of DNA synthesis, preventing proliferation of irreversible DNA damage in future cells (Bertoli et al., 2013).

Differential expression of transcripts involved in mRNA splicing (pathways: spliceosomal complex assembly, RNA splicing; hub transcripts: *zmat5, cwc27, prpf31*) highlight modified post-transcriptional RNA processes occurring as temperatures increased. Post-transcriptional modifications by pre-mRNA splicing can provide increased transcriptional plasticity, with the potential for enhanced acclimatory capacity when faced with environmental change (Mastrangelo et al., 2012; Steward et al., 2022; Thorstensen et al., 2022b). However, ultimately as temperatures reached 24^°^C, these processes were downregulated across populations, demonstrating the impacts that increasing temperatures have on transcriptional plasticity, reducing the capacity for physiological plasticity through alternative splicing mechanisms as stressors mount.

As temperatures increased, there were modifications to protein synthesis and turnover processes (pathway: protein catabolic process, protein ubiquitination, protein polyubiquitination; hub transcripts: *rps27a, cops5, hap1, rnf8, usp13, fbxo45, smurf2, rplp0, bop1, eif3d*) key to translation, ubiquitination, and amino acid recycling. Interestingly, *rps27a,* with dual roles providing both ubiquitin and ribosome structural components (Klinge et al., 2012; Luo, et al., 2022), was at the center of the network with the largest number of protein-protein interactions and betweenness score, emphasizing its role within the regulatory network in response to thermal stress. These protein synthesis and recycling mechanisms are key to modifying the organism’s existing phenotype as environments shift, and without their proper function may leave the organism vulnerable to the effects of compounding stressors (Davies et al., 2016).

Increased temperatures also modulated chaperone activity and protein folding processes (pathways: protein folding, regulation of protein stability, response to heat; hub transcripts: *hsc70, hspa8, hyou1, dnajc2, cct4, unc45b, naca*) in the face of thermal stress and reactive oxygen species formation, consistent with a broad literature of molecular responses to thermal stress (Place and Hoffman, 2001; Jackson and Hewitt 2016; Vallin and Grantham, 2019; Penny et al, 2023). Modulation of these chaperone proteins, paired with an observed decrease in metabolic rate for lake sturgeon in temperatures of 24^°^C from previous research (Bugg et al., 2020), suggest that there was a transcriptional modification of heat shock protein production under thermal stress. Transcriptionally modification of heat shock protein production and decreased mitochondrial oxygen consumption (Bugg et al., 2020) could aid to suppress the formation of reactive oxygen species as a mechanism to stabilize the existing cellular proteome to enhance long-term fitness and survival (Storey and Storey, 2011).

Finally, elevated temperatures altered immune system processes (pathways: inflammation, response to virus, immune system process, adaptive immune response, cytokine-mediated signaling pathway, lymphocyte differentiation; hub transcripts: *traf6, b2m, banf1, rsad2, egr1, jak1, prkcbb*). Pathways and related hub genes indicate that there were altered innate and adaptive immune markers associated with viral infection suggesting immune compromise and increased susceptibility to pathogens as temperatures increased, with a potential threshold of 20^°^C for immunocompetence (Choi, 2005; Li et al., 2016; Wei et al., 2017; Ebrahimi, 2018). Thus, future increases in temperature will not only have direct effects on the lake sturgeon’s thermally responsive mechanisms, but likely indirect effects on pathogen tolerance and survival as combining stressors mount (Dittmar et al., 2014; Alfonso et al., 2020).

### Mitochondrial Limits Under Thermal Stress

Based on transcriptional evidence, mitochondrial function is likely inhibited across both populations of lake sturgeon as temperatures increased to 24^°^C. Many mitochondrial processes were downregulated in response to initial thermal stress at 20^°^C and continued to decline as temperatures increased. Transcriptional markers of dysfunction and resulting damages were apparent at 20°C in northern Burntwood River sturgeon and were more extreme when compared to southern Winnipeg River sturgeon, highlighting potential population-specific mitochondrial capacity. Additionally, a highly differentially expressed hub transcript (*gfm1*) key to mitochondrial translation and oxidative phosphorylation (Berenguer et al., 2021) was altered as temperatures increased. Aside from the direct effects of decreased ATP production and increased reactive oxygen species damage, a decline in mitochondrial function also has implications for stress signaling, viral immune response, and formation of heme as well as iron cluster assemblies required for oxygen transport and numerous cellular processes (Tovar et al., 2003; Ajioka et al., 2006; Lin et al., 2006; Qin et al., 2010; Karnkowska et al., 2016; Hernansanz-Agustin et al., 2020; Braymer et al., 2021). The impairment of these widespread mitochondrial processes in developing lake sturgeon may inhibit the function of the mitochondria itself, related cellular systems, and ultimately gill function, with the potential for population-specific thresholds to impact whole-organism fitness during changes in environmental temperatures (Iftikar and Hickey, 2013; Jeffries et al., 2018).

### Study Limitations

As lake sturgeon are an octoploid species, transcriptomic analysis can be vulnerable to the effects of gene duplication, resulting homeologs, and their effect on the interpretation of resulting sequence data (Ogden et al., 2013; Limborg et al., 2016). To mitigate this effect, we selected a tissue-specific, *de novo*, sequencing approach to help isolate different gene isoforms. Additionally, we selected a transcriptome assembly program, Trinity, that has been shown to have success with the accurate assembly of polyploid transcriptomes (Grabherr et al., 2011; Voshall & Moriyama, 2020). With respect to SNP calling using the transcriptome, we used a splice-aware read mapping program (STAR; Dobin et al., 2013) and addressed paralogous loci with heterozygosity and read depth ratio deviations (McKinney et al., 2017), in addition to standard quality control and linkage disequilibrium pruning steps. Nevertheless, the genetic differentiation presented in the present study is biased with family structure by our experimental design (studied individuals represent F_1_ crosses of wild-caught adults), and therefore are not representative of differentiation between the two studied populations overall (Kjartanson et al., 2023). For instance, the PCA with SNPs may not accurately reflect patterns of genetic variation in F0 wild individuals. Nor are these genetic data appropriate for studying heritability or additive genetic effects because of the difficulties of ploidy and SNPs with mRNA data. Instead, the genetic data gathered from fish in each population provides context for the observed transcriptional plasticity, which is at least partially attributable to genetic differences between the individuals used in the study.

While the octoploid lake sturgeon genome can complicate transcriptomic analysis (Ogden et al., 2013; Limborg et al., 2016), the consistency in responses to temperature across the populations, and with other literature, lends credence to the reliability of the interpretation of sequencing data. Moreover, the thermal resolution of our study (4^°^C windows) may make the exact detection of thermal thresholds difficult, especially if thresholds are non-linear across populations. Further, this work only represents the findings from two latitudinally distributed populations. Future research should expand these findings across the full latitudinal distribution of lake sturgeon, or across other latitudinally distributed sturgeon species.

### Ecological Relevance

In the current study, temperatures of 20^°^C and above limited the transcriptional plasticity of developing lake sturgeon, which would ultimately diminish their plasticity to other stressors. As temperatures continue to increase in the wild, the combined effects of thermal, oxidative, and pathogen related stressors will likely limit the survival of lake sturgeon populations within Manitoba, Canada, with increased severity for northern sub-Arctic populations throughout the province (Bugg et al., 2020). The population designation units now group lake sturgeon inhabiting northern regions with those from several other populations in Manitoba to form a larger designation unit (COSEWIC 2006; COSEWIC 2017). However, new evidence suggesting large discrepancies in transcriptomic, physiological, and genetic differences between populations, as well as their natural thermal environments and tolerance thresholds (McDougall et al., 2017; Bugg et al., 2020; Bugg et al., 2021; Kjartanson et al., 2023) indicate that they are differentially susceptible to increasing temperatures and require distinct management strategies for conservation. While the experimental design used precludes analyses of heritability and additive effects of genetic variation underlying the observed transcriptional plasticity, these results are nevertheless valuable for their physiological and population-level implications.

Currently, temperatures in the Winnipeg River exceed 20^°^C for approximately 50 days per year, while this temperature threshold is projected to be exceeded during June, July, and August in the Burntwood River in the coming decades (Manitoba Hydro, 2015; Vincent et al., 2015; Zhang et al., 2019; Bugg et al., 2020). These seasonal increases in temperature occur when newly hatched sturgeon are in their first few months of life, which are the same life stages examined in the current study, and could exacerbate already high rates of mortality during this period (Bugg et al., 2020; Bugg et al., 2021). As temperatures increase transcriptional modification occurs, but many key mechanisms across both populations are downregulated by 24^°^C. Additionally, endemic viral pathogens are persistent in both northern and southern populations of lake sturgeon throughout the province, with higher abundance and probability of detection in younger fish (Clouthier et al., 2013; Clouthier et al., 2020), and as temperatures increase. With the present experimental evidence, we suggest that if river temperatures continue to increase as projected (Manitoba Hydro, 2015) and there are not sufficient thermal refugia, these physiological limitations will likely impede future lake sturgeon recruitment, especially in the most northern populations. Therefore, increasing temperatures may influence recruitment, movement patterns, and distributions of lake sturgeon within Manitoba, as sturgeon shift to areas of thermal refuge to preserve energy and persist in the face of environmental change (Moore et al., 2020; Robertson et al., 2021). However, even with sufficient refugia, fish do not always travel to habitat with increased suitability due to life history constraints (Sutton et al., 2007; White et al., 2019). If lake sturgeon are unable to redistribute themselves in areas of thermal refuge, evidence suggests that the impacts of increasing environmental temperature may be more severe, with impacts to population-level processes and survival based on observed transcriptional plasticity and physiological responses when exposed to elevated temperatures.

## Data Accessibility

The data used in this study is available in the included supplementary files (1-8) and the raw sequencing reads are available at the National Center for Biotechnology Information Sequence Read Archive with the accession number PRJNA815828. The Trinity assembly and Trinotate annotation file are available via figshare (https://doi.org/10.6084/m9.figshare.19209753.v1).

## Author Contributions

WSB, WGA, and KMJ designed the research. WSB analyzed the count data with assistance from KM and MJT. MJT analyzed the single nucleotide polymorphism data. WSB wrote the first draft of the manuscript and all authors provided comments on the final version.

## Acknowledgements

The authors thank North/South Consultants Inc. and Manitoba Hydro for their assistance in the capture of spawning adults and collection of gametes from these wild sturgeon populations, Evelien de Greef for her support with the map (Figure 1), as well as Madison Earhart for valuable discussions and sturgeon illustrations (Figure 2 and Figure 3A). We would also like to thank the staff at the University of Manitoba Animal Holding Facility and numerous Anderson and Jeffries Laboratory members for assistance in the care and maintenance of fish. This work was conducted at the University of Manitoba located on original lands of the Anishinaabeg, Cree, Oji-Cree, Dakota and Dene peoples, and on the homeland of the Métis Nation. We recognize that water supplied for our fish at the University of Manitoba was sourced from the Shoal Lake 40 First Nation. Many bioinformatics steps were enabled by our chance to use Digital Research Alliance of Canada computing clusters. Funding for this study was provided by NSERC/Manitoba Hydro Industrial Research Chair awarded to W.G.A. and NSERC Discovery Grants (grant numbers 05348 and 05479) awarded to W.G.A. and K.M.J., respectively. W.S.B. was supported by a University of Manitoba Graduate Fellowship as well as the International Graduate Student Scholarship.

## Supplemental Figures

**Supplemental Figure 1.**
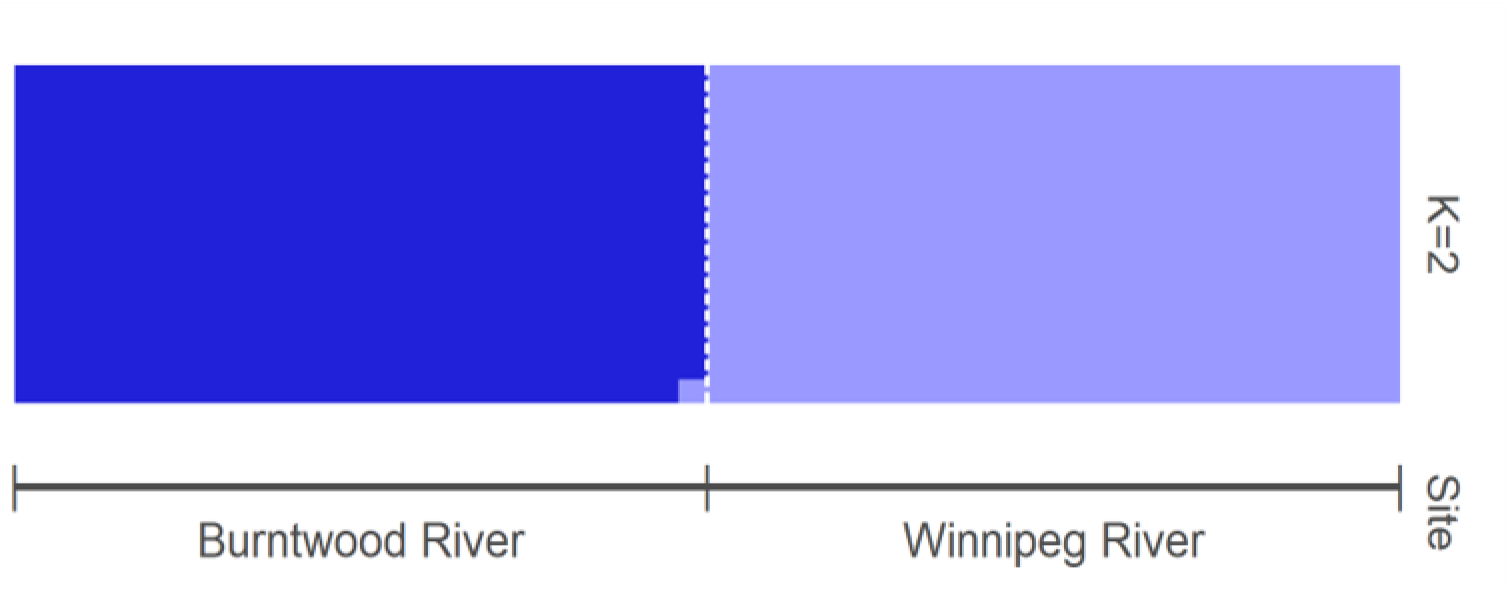
Admixture plot illustration the distribution of SNPs between northern Burntwood River and Southern Winnipeg River populations of lake sturgeon, *Acipenser fulvescens*, within Manitoba, Canada.

**Supplementary Figure 2.**
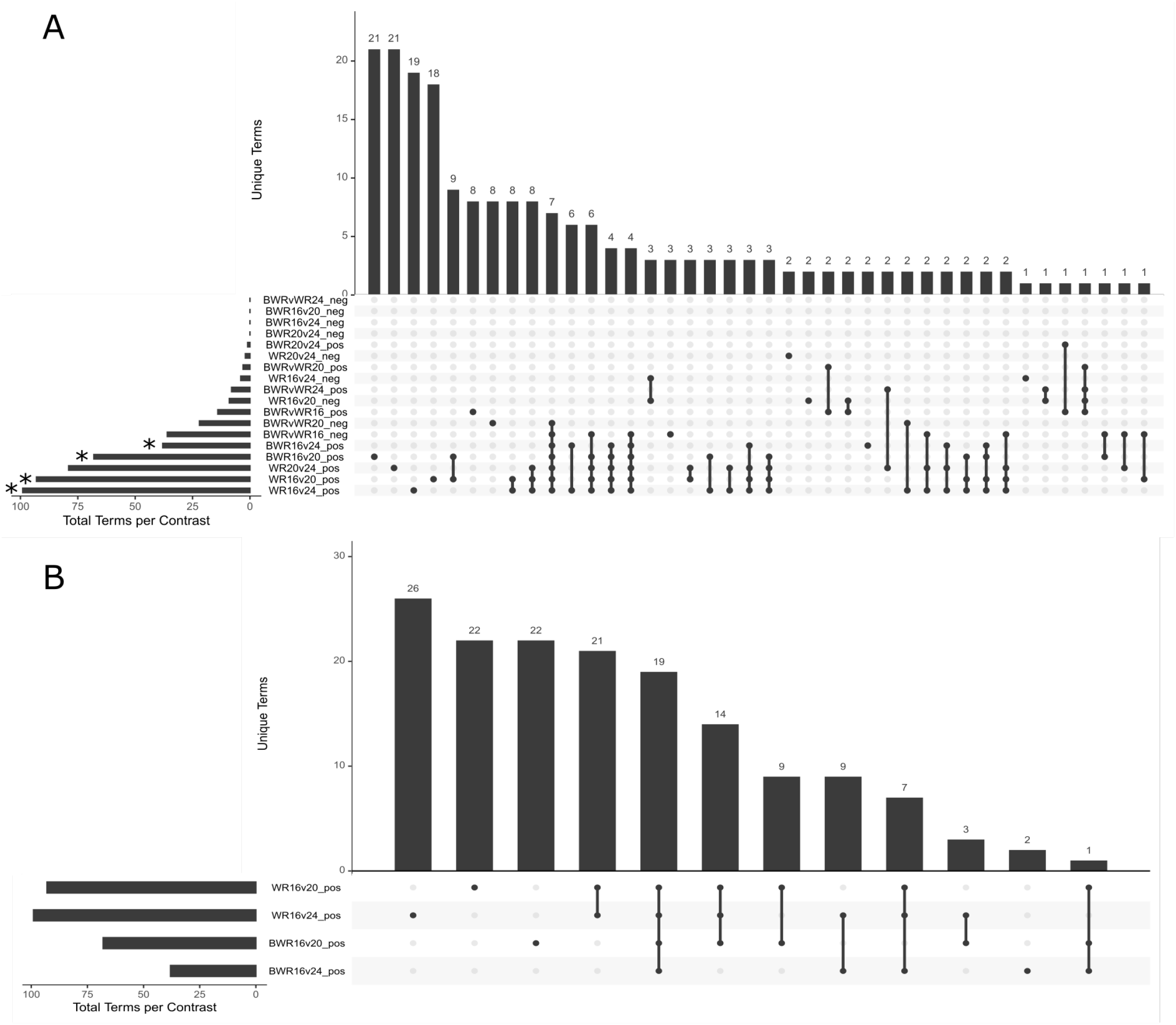
GO (gene ontology) term distribution illustrated using UpSet plots demonstrating shared and population-specific terms across A) all split-contrasts and B) split contrasts with terms decreasing as environmental temperatures increase. *’s on split-contrast in A indicate the contrasts which were included in B. These data represent change in the transcriptional mechanisms in the gill of developing lake sturgeon from both a northern Burntwood River (BWR) and southern Winnipeg River (WR) population of lake sturgeon (*Acipenser fulvescens*), within Manitoba, Canada, acclimated to temperatures 16^°^C, 20^°^C, and 24^°^C for 30 days in early development, measured using RNAseq. Individual dots indicate processes unique to that contrast, while a line connecting multiple dots indicate processes that are shared between the contrast connected by the line. Split-contrasts are oriented BWR-WR at the interpopulation level and lower temperature-higher temperature across acclimation treatments so that “neg” indicates upregulation at higher temperatures, and “pos” indicates downregulation at higher temperatures (e.g. WR16v20_pos includes terms that were downregulated in the WR as temperatures increased). Each population and acclimation treatment is represented by 6 individual gill samples (n = 6).

**Supplementary Figure 3.**
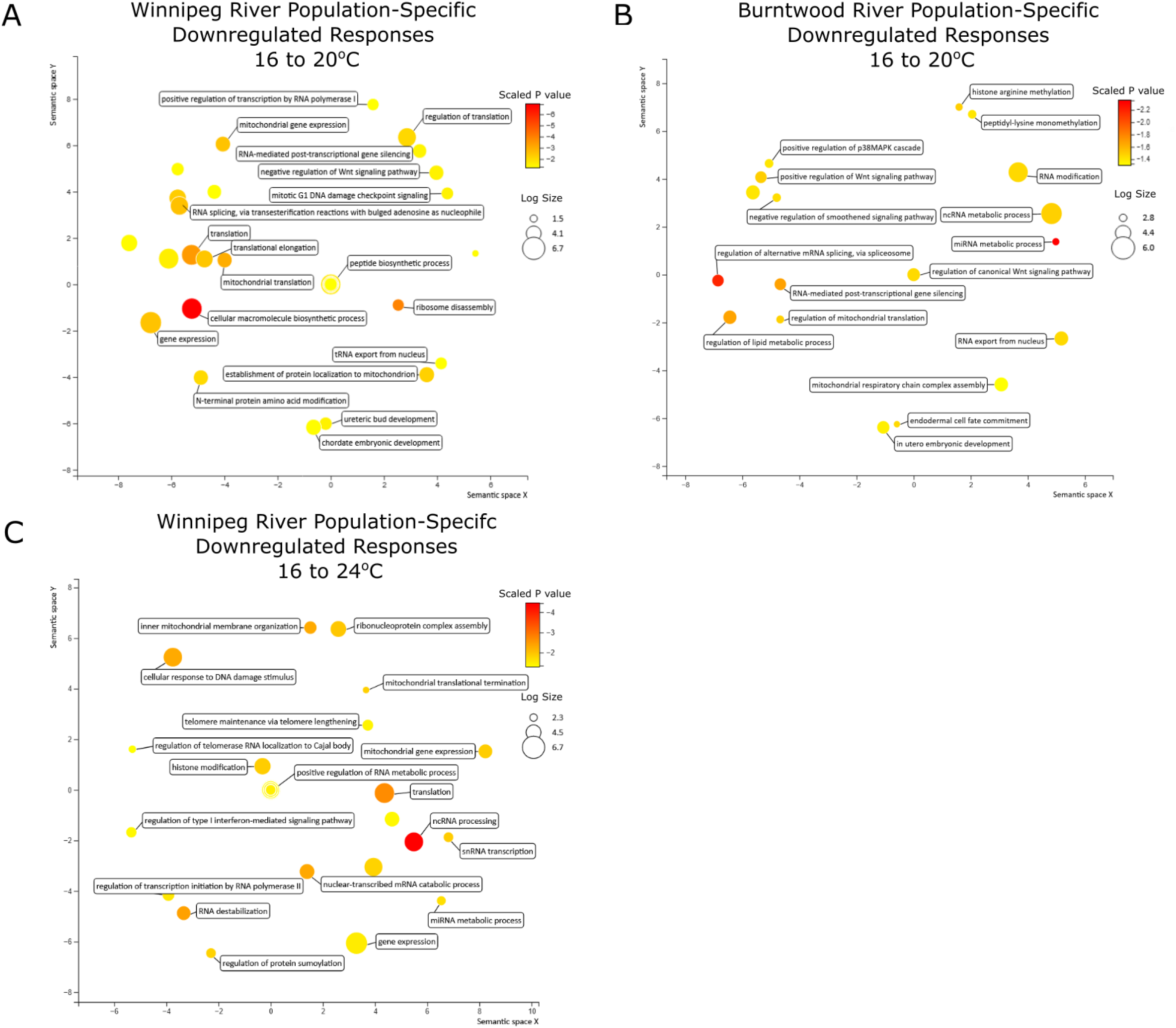
REVIGO (Reduce + Visualize Gene Ontology) plots of GO (gene ontology) population-specific biological processes terms for the Winnipeg River and Burntwood River populations of developing lake sturgeon (*Acipenser fulvescens*) within Manitoba, Canada, as acclimation temperatures increase during a 30-day acclimation, measured using RNAseq. Plots represent A) Winnipeg River population-specific responses downregulated from 16 to 20^°^C B) Burntwood River population-specific responses downregulated from 16 to 20^°^C) Winnipeg River population-specific responses decreased from 16 to 24°C. Bubble color indicates the significance of the adjusted *p* value after applying a log_10_ scale (darker color is a more significant term), while bubble size indicates the frequency of a GO term GO annotation database (larger bubble is a more common GO term in the data set).

## Notes

https://doi.org/10.6084/m9.figshare.19209753.v1

### Competing Interest Statement

The authors have declared no competing interest.

### Summary of Updates

This version of the manuscript has been revised to update the analysis as there was an error found in the metadata upon initial review. All analysis has been re-conducted with the correct metadata and the discussion has been rewritten to reflect the new results. Further, SNP level analysis has been added to the manuscript.

